# GeMSTONE: Orchestrated Prioritization of Human Germline Mutations in the Cloud

**DOI:** 10.1101/052001

**Authors:** Siwei Chen, Juan Felipe Beltrán, Xiaomu Wei, Steven Lipkin, Clara Esteban-Jurado, Sebastià Franch-Expósito, Sergi Castellví-Bel, Haiyuan Yu

**Author notes:** The authors wish it to be known that, in their opinion, the first 2 authors should be regarded as joint First Authors.

## Abstract

Integrative analysis of whole-genome/exome-sequencing data has been challenging, especially for the non-programming research community, as it requires leveraging an inordinate number of computational tools. Even computational biologists find it unexpectedly difficult to reproduce results from others or optimize their own strategies in an end-to-end workflow. We introduce **Ge**rmline **M**utation **S**coring **T**ool f**O**r **N**ext-generation s**E**quencing data (GeMSTONE), a cloud-based variant prioritization tool with high-level customization and a comprehensive collection of bioinformatics tools and data libraries (http://gemstone.yulab.org/). GeMSTONE generates and readily accepts a sharable “recipe” file for each run to either replicate existing results or analyze new data with identical parameters.

X.W., S.L. and H.Y. conceived of the GeMSTONE concept. S.C. designed and implemented the GeMSTONE pipeline with supervision and input from H.Y. and X.W. J.F.B. designed and implemented the GeMSTONE web interface. C.E.J., S.F.E. and S.C.B. provided CRC study datasets and the original study design. J.F.B., S.C. and H.Y. wrote the manuscript. All authors reviewed and approved the final manuscript as submitted.

The authors declare no competing financial interests.

---

Next-generation sequencing (NGS) has greatly reduced the costs of obtaining genomic data for increasingly large sample sizes^1^, facilitating discovery of causal gene and mutation candidates for various disorders^2^ and providing sizable genetic variant datasets^3^. As a result, the process of filtering, annotating, and prioritizing variants from large-scale studies has grown in complexity and computational burden. It has quickly become difficult to organize, maintain, and standardize the constituent parts of thorough variant analysis, which increases the time and dime investment for less computationally oriented biologists and labs. Even now, some researchers write custom scripts to analyze genome-wide genetic variation data step-by-step^4–6^, a task so often undertaken that standardization is long overdue. Moreover, without a standard set of tools and conventions, analytic protocols can grow vague and hard to replicate^3^, even within the same lab.

We present GeMSTONE, a cloud-based variant prioritization framework that leverages 7 popular bioinformatics suites (Vt^7^, VCFtools^8^, BCFtools^9^, SnpEff^10^, GEMINI^11^, dbNSFP^12^, and PLINK/SEQ^13^) and connects 46 meta-information and prediction resources (**Table 1**) to provide a smooth, customizable workflow for variant analysis. GeMSTONE provides a code-free portal for filtering, annotation, and prioritization, which not only helps standardize genetic variation analyses, but also offers the means to easily replicate and share computational protocols. From a user’s perspective, GeMSTONE is a reliable one-stop shop for variant analysis where they can find a collection of tools spanning a broad range of applications through an intuitive, unified user interface subsuming all general-purpose workflows from comparable toolkits (**Figure 1**).

**Table 1.**
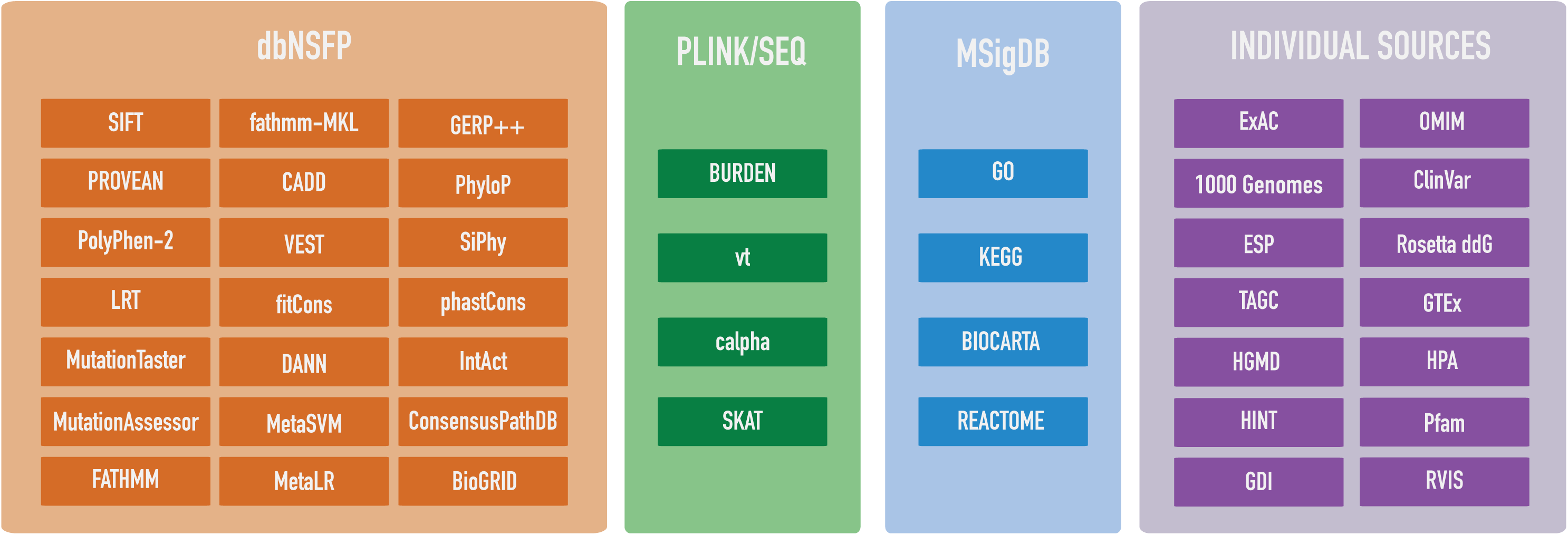
**Data resources and bioinformatics tools built in GeMSTONE**

**Figure 1.**
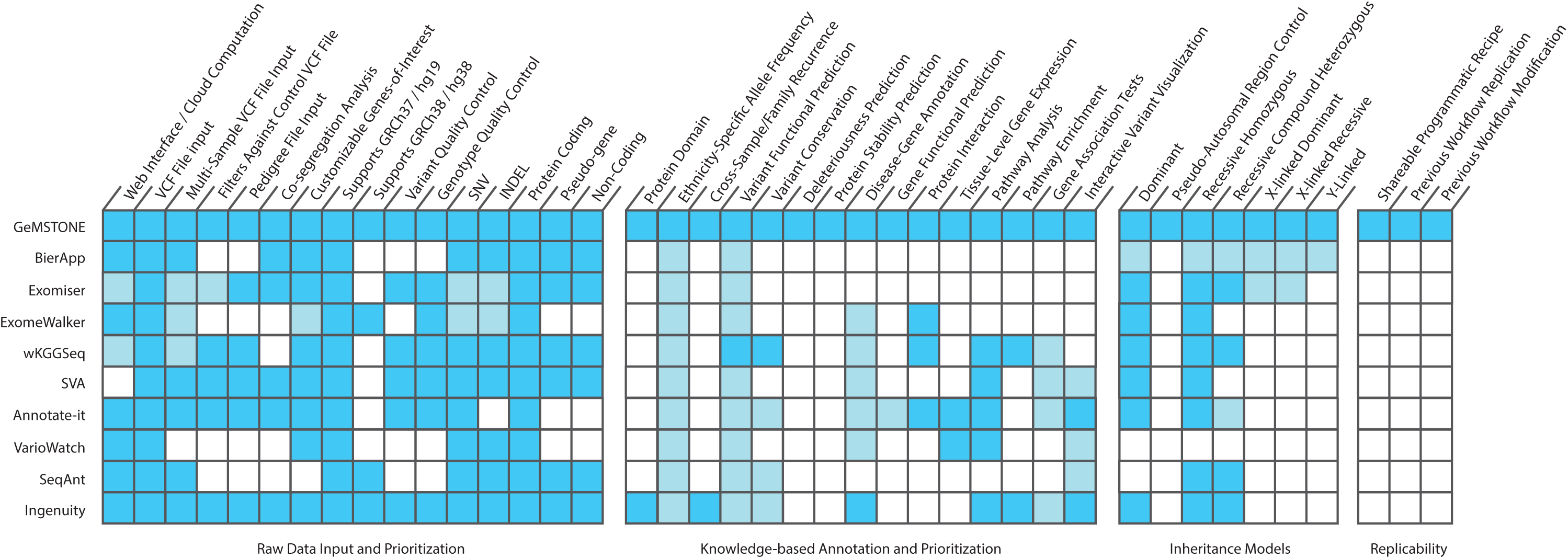
Heatmap comparison of GeMSTONE and other variant analysis web. This heatmap compares different aspects of GeMSTONE’s functionality and other web tools with similar objectives. Each row represents a different tool, while each column represents a specific feature. Dark blue indicates that a tool has similar capacity for a specific function while light blue indicates that a tool has a similar feature but with less powerful functionality than GeMSTONE (see Supplementary Note 1).

Users of the GeMSTONE web portal can customize their analyses of genomic data from Variant Call Format (VCF) files by using a range of tool classes, including basic raw-data filters on genotype quality and variant consequences, variant/gene-based filters on functional predictions to more comprehensive annotations on protein-protein interaction network and biological pathways across vast knowledge-based databases. The user may also choose to include supplementary files, such as a pedigree (PED) file for co-segregation analysis with a selected inheritance model, or a list of genes for customized annotation. These kinds of workflow decisions are all part of a highly customizable filtering, annotation, and prioritization pipeline (**Figure 2**). Once a user has set their desired parameters for a run, it is scheduled for processing on a protected server, alleviating the user’s burden to update software, parse data libraries, store large derivative files, and dedicate processing time. Once their job is finished, the user can log into the GeMSTONE portal to selectively download step-by-step snapshots of their workflow, interactively visualize their variant statistics (**Supplementary 4**), and download a recipe (JSON) file that can be uploaded to share, replicate, or modify the same workflow.

**Figure 2.**
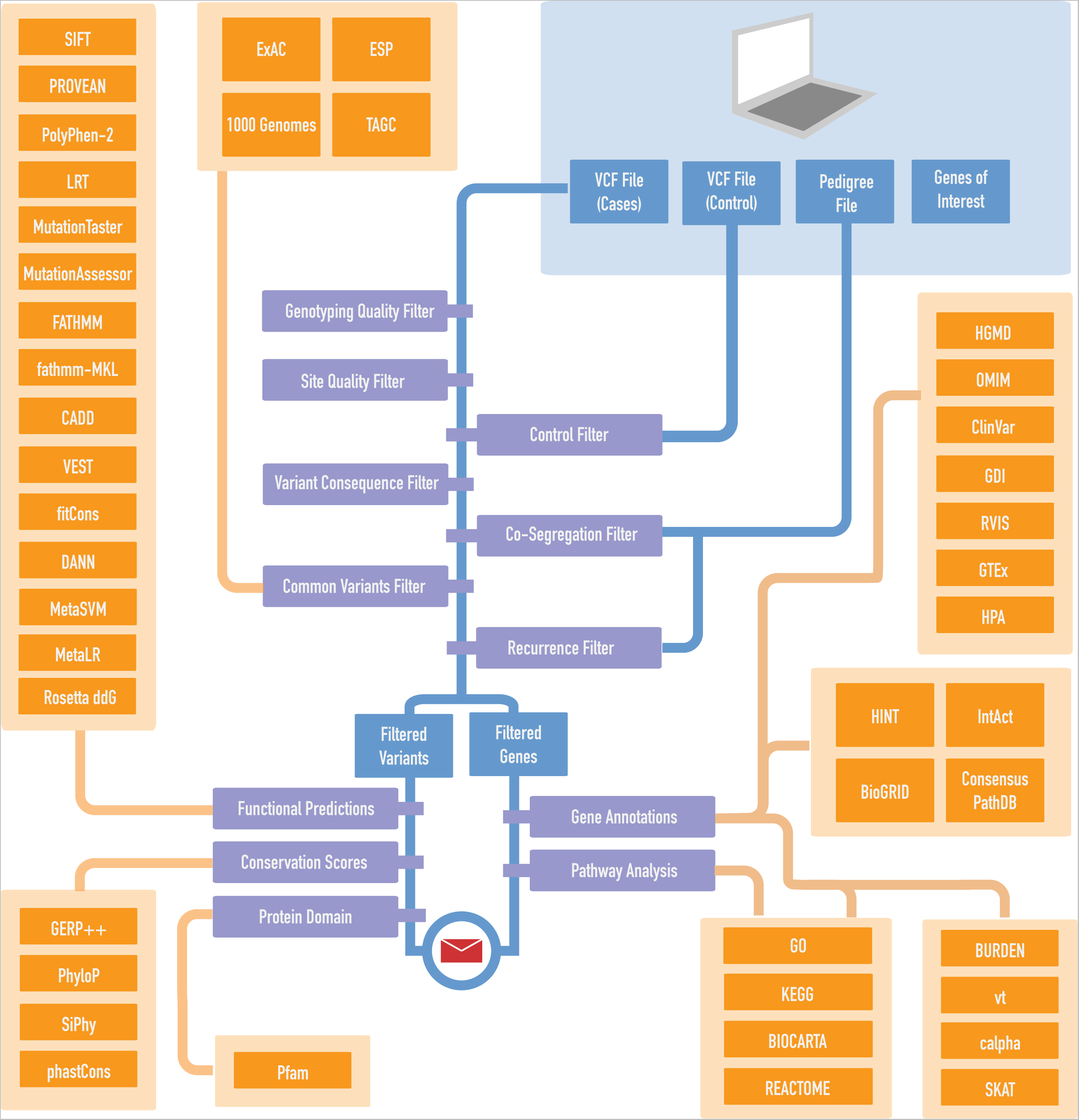
GeMSTONE pipeline overview. The schematic represents the GeMSTONE’s central analysis pipeline. The fundamental backbone filter cascade can be seen in blue, prioritizing rare and putatively damaging variants, as well as genes of high sequencing quality. Different libraries are grouped in orange, participating in annotation or filtering steps throughout the workflow as indicated.

A major design goal in the development of GeMSTONE is the ability to maximize customization for studies with very specific methodology without overwhelming users who want to execute a quick, exploratory workflow. For instance, GeMSTONE’s variant deleteriousness prediction and protein stability calculation toolset boasts 23 algorithms and data libraries (**Figure 2**). Users can choose to annotate using any of these predictors, and even customize the ‘deleteriousness range’ of each individual predictor. A global deleteriousness filter is available, which allows users to set a threshold on the number of selected deleteriousness predictors needed in order for a variant to pass the filter. This set of filters is useful in that it allows the user to adjust the stringency of defined algorithms and to balance any inconsistency among different predictions. This collection of deleteriousness prediction tools allows users to choose which algorithms they use solely based on their relative merit as metrics, rather than the programming investment that it would take to install or query them.

Most options within the GeMSTONE workflow can serve a double purpose as either filters or simply as annotations. For the global deleteriousness filter mentioned above, the count of deleterious predictions and their individual scores will be annotated next to each variant, providing information that can be used for variant prioritization without being part of any filter. Another example of flexible library use is the allele frequency database menu. Users can choose to annotate all variants with allele frequencies from 4 different databases (ExAC^14^, 1000 Genomes^15^, ESP6500^16^, and TAGC^17^) and/or filter by allele frequency in relation to any sub-population within them in an sample-specific manner. We also provide the option to combine information across libraries—for instance, allowing for known disease gene annotation on candidates to be supplemented with their interaction partners as reported in other databases. These kinds of tools (**Table 1**) have never been collected, maintained or connected in a single resource before (**Figure 1**). By collecting and connecting this suite of tools GeMSTONE seeks to maximize workflow utility for many different use-cases—whether it is stringent filtering to eliminate noise in massive sequencing datasets, or data exploration in small familial studies.

In particular consideration of relatively large sample sizes, GeMSTONE allows users to set recurrence filters, a minimum and maximum number of times variants must be shared by sporadic samples and/or families to pass the filter. This filter was implemented following user tests, as we found this option—though seldom described—was very much desired when dealing with datasets containing large number of sporadic samples to prioritize high risk disease-associated mutations while exclude potential sequencing artifacts. This process of user-driven development by which GeMSTONE morphs to the community’s needs is the key behind GeMSTONE’s ability to grow as a knowledge bank with a robust and updated set of functionalities. Small but important filtering steps like these, now explicitly documented in GeMSTONE summary and recipe files, can become part of study replication without any issue.

By maintaining an updated set of bioinformatics tools for variant analysis, GeMSTONE provides a decreased barrier to entry for less computationally oriented research groups, and a central hub for researchers working on genomic variation studies. The options offered by the web interface serve as a way for users to explore and learn about new tools and data sources, while also providing developers with an overview of the current variant analysis landscape in order to fill any gaps in the current tool-space. New tools can be easily added to GeMSTONE, and presented to the community with less platform-specific barriers.

As an example of a GeMSTONE use case, we replicated a published analysis of rare pathogenic variants in new predisposition genes for familial colorectal cancer (CRC)^4^. The original workflow of the study included steps such as deleteriousness prediction cutoffs across different algorithms, as well as pathway analysis of associated genes (**Supplementary 1**). We performed the entirety of the authors’ analysis through a single run of GeMSTONE. In the same vein, we prioritized variants from a whole exome sequencing dataset containing 247 amyotrophic lateral sclerosis (ALS) patients and 268 healthy samples as controls (**Supplementary 2**). We analyzed the intermediate VCF files at each step of the pipeline, showing that more stringent filters produced a set of variants that are much more localized to protein-protein interaction interfaces derived from co-crystal structures (**Supplementary 3**). Variants from the raw data have a random distribution throughout the protein (odds ratio = 1.04), whereas over one-third of GeMSTONE-filtered variants are located on protein-protein interaction interfaces, a significant enrichment with respect to the relative size of interfaces compared to whole proteins (odds ratio = 2.5, *P* < 10^−5^ with a *Z*-test). We conducted the same enrichment analysis on the CRC replication set and found the same trend to be true (**Supplementary 3**). The ability to track these trends through different filtering steps is a great way to both introspect and debug filtering decisions. More importantly, the GeMSTONE workflow allows biologists to complete these kinds of scripting-heavy protocols without writing a single line of code.

GeMSTONE opens up the door for accessible, collaborative, replicable, and holistic analysis of genetic variants. First, it seamlessly knits together filters and annotations through different tools with either stringent, study-specific parameters or general best-practice settings. Second, it eliminates the time and space burdens associated with modern variant analysis tools, saving users dozens of gigabytes of potential disk-space per run for the same workflow on a medium-sized dataset. Third, it significantly lowers the barrier to entry for traditional biologists by eliminating the installation and scripting sinkholes that may dissuade researchers from pursuing large-scale analysis or trying new tools. Fourth, it provides a readable, shareable log—both programmatic and human—to allow other researchers to understand and replicate study results given the same starting data. Finally, GeMSTONE encourages growth of the genomics research community by maintaining and updating a bank of best-practice bioinformatics methods and tools.

## Acknowledgements

IDIBAPS: We are sincerely grateful to the patients and their families for their participation. We are really thankful to the Centre Nacional d’Anàlisi Genòmica and the Biobank of Hospital Clínic–IDIBAPS, Barcelona, for technical help. The work was carried out (in part) at the Esther Koplowitz Centre, Barcelona. CEJ and SFE are supported by a contract from CIBERehd. CIBERehd is funded by the Instituto de Salud Carlos III. This work was supported by grants from Fondo de Investigación Sanitaria/FEDER (14/00173), Fundación Científica de la Asociación Española contra el Cáncer (GCB13131592CAST), COST Action BM1206, Beca Grupo de Trabajo “Oncología” AEG (Asociación Española de Gastroenterología), and Agència de Gestió d’Ajuts Universitaris i de Recerca (Generalitat de Catalunya, 2014SGR255).

